# Stage-sensitive potential of isolated rabbit ICM to differentiate into extraembryonic lineages

**DOI:** 10.1101/2023.12.19.571672

**Authors:** Katarzyna Filimonow, Anna Chołoniewska, Jan Chołoniewski, Zofia E. Madeja, Katarzyna Barłowska, Joanna Grabarek, Berenika Plusa, Anna Piliszek

## Abstract

In the course of mammalian development initial state of totipotency must be lost to allow acquisition of specific cell fates. The first differentiation event results in the formation of trophectoderm (TE) and the inner cell mass (ICM). In the mouse embryo the cell fate of these two compartments is set quickly after formation of a blastocyst. However, recent reports suggest that plasticity of these two lineages might be extended in species other than the mouse. Here we investigated how the cellular plasticity of early mammalian embryos relates to developmental time scale and changes in gene expression using rabbit isolated ICMs. We studied the dynamics of rabbit blastocyst formation using time-lapse imaging and identified GATA3 as an early marker of rabbit TE and CDX2 as a marker of fully formed TE. We then analysed developmental potential of rabbit ICMs isolated by immunosurgery and subsequently cultured *in vitro*. ICMs originating from early- to mid-blastocyst stage embryos are able to re-form a blastocyst-like structure, with a functional TE, and an ICM containing both SOX2-positive epiblast cells and SOX17-positive primitive endoderm cells. We further observed that rabbit ICMs isolated from later blastocyst stages lose the ability for TE specification, instead forming a halo-like cavity with an outer layer of SOX17-positive cells. Our data indicate that in mammalian embryos the potential for TE differentiation gives way to formation of a different type of extraembryonic epithelial layer, suggesting potential common mechanism of pluripotency restriction between eutherian mammals.

## INTRODUCTION

The early stages of embryonic development of eutherian mammals are devoted to differentiation of the first cell lineages – pluripotent epiblast (Epi), and extraembryonic primitive endoderm (PrE) and trophectoderm (TE). TE is the first cell lineage specified during mammalian ontogenesis. Proper differentiation of TE is a prerequisite for supporting the pregnancy, as TE is responsible for embryo implantation in the uterus and forms the embryonic part of the placenta (reviewed in (Hemberger et al., 2020)). Placental related disorders affect around one-third of human pregnancies (Jauniaux et al., 2006), and TE quality and cell number can be used as an effective predictor of successful pregnancy in human assisted reproductive technologies (Ahlström et al., 2011; Ebner et al., 2016). Despite a significant knowledge about TE lineage specification in the mouse, data concerning this first differentiation event in other mammalian species, including human and domestic animals, are still limited. Importantly, a growing body of evidence indicates that notable differences exist in early lineage specification between mouse and other eutherian mammals. Embryos of human or primate origin are not easily available for research due to the ethical concerns and the lack of easily accessible material. In addition, many experimental approaches are not applicable when using non-rodent systems and embryological studies in both primates and large domestic animals mainly rely on *in vitro* embryo production (IVP). Even the best IVP conditions are suboptimal when compared to *in vivo* environment, and several reports show a significant influence of *in vitro* culture conditions on lineage allocation and lineage-specific gene expression in mammalian preimplantation embryos (Choi et al., 2015; Henderson et al., 2014; Saenz-de-Juano et al., 2013). Therefore, it is critical that important developmental processes, such as TE specification, are studied in both *in vitro* and *in vivo* derived embryos.

In the mouse embryos, specification of the TE lineage from the outside cells of the morula is regulated by the expression of several transcription factors, including Caudal-related homeodomain transcription factor (CDX2) (Strumpf et al., 2005) and GATA-binding protein 3 (GATA3) (Home et al., 2009; Ralston et al., 2010). *Cdx2* expression in the mouse is already detected prior to cavitation, and at the blastocyst stage it becomes restricted to TE cells (Strumpf et al., 2005). CDX2 is also a specific TE marker in non-rodent mammals, including human, pig, cattle and rabbit (Berg et al., 2011; Bou et al., 2017; Chen et al., 2012a; Goissis and Cibelli, 2014; Kuijk et al., 2008; Madeja et al., 2013; Sakurai et al., 2016). However, in contrast to the mouse, in most of these species CDX2 expression has not been confirmed prior to cavitation. GATA3 is also associated with TE fate specification in the mouse (Home et al., 2009; Ralston et al., 2010). GATA3 is co-expressed with CDX2 in mouse TE, and it is capable of inducing TE specification in mouse embryonic stem cells (ESCs) (Ralston et al., 2010). GATA3 has been also shown to be a specific TE marker in the horse (Iqbal et al., 2014), cattle (Gerri et al., 2020; Ozawa et al., 2012; Smith et al., 2010) and human embryos (Gerri et al., 2020; Guo et al., 2021).

At the initiation of embryonic development, individual cells (blastomeres) are totipotent and equal in their developmental potential (Mintz, 1965; Tam and Rossant, 2003; Tarkowski, 1959; Tarkowski and Wróblewska, 1967). This early totipotency has to gradually give way to the differentiation of the specific cell lineages and tissues, in order for the embryo to develop correctly. In the mouse, TE versus inner cell mass (ICM) cell fate becomes mostly restricted around the time of cavitation (Gardner, 1983; Nichols and Gardner, 1984; Suwińska et al., 2008). In contrast, human and bovine embryos retain high plasticity up to day 6 of development (Guo et al., 2021; Kohri et al., 2019). In cattle, ICMs isolated from day 6 blastocysts are able to regenerate the TE layer, forming blastocyst-like structures that support full-term development (Kohri et al., 2019), whereas human naïve epiblast can regenerate TE cells (Guo et al., 2021). Currently, it still remains to be resolved whether a shorter period of ICM plasticity in the mouse represents a significant deviation from other mammals.

To further understand these interspecific differences in TE versus ICM lineage specification during mammalian development, we looked for the model organism that allows for analysis of *in vivo* fertilised embryos as well as experimental manipulation. Lagomorphs (including rabbits) have been reported to be genetically closer to Primates than Rodents are (Allard et al., 1996; Graur et al., 1996). Rabbit *in vivo* fertilised embryos are easily available for study, which allows avoiding any potential anomalies in development induced by *in vitro* fertilisation procedure. Here, we present an analysis of TE-associated transcription factors expression and their spatio-temporal localization at the consecutive stages of rabbit blastocyst development *in vivo*, as well as time-lapse imaging-based examination of rabbit TE morphogenesis. We further investigate rabbit ICM commitment and potency by analyzing the regenerative potential of isolated rabbit ICMs. Our study shows that although TE-associated transcription factors are conserved among mammals, their regulation might significantly differ, especially between rodents and non-rodent species. We also reveal notable differences in the early lineage differentiation potential between species, evidenced by the delayed differentiation of rabbit ICM compared to the mouse.

## MATERIALS AND METHODS

### Animals

Rabbits (*Oryctolagus cuniculus*, Popielno breed) were maintained under a 14-h light/10-h dark cycle in the animal facilities of The Institute of Genetics and Animal Biotechnology of the Polish Academy of Sciences (IGAB PAS) according to the national and institutional guidelines. Experimental procedures were approved by the Second Local Ethics Committee, Warsaw, Poland (permission number: WAW2/183/2018 and 31/2012).

### Embryo collection and culture

Embryos were derived following natural matings, surgically from females under general anaesthesia or from dissected reproductive tract of sacrificed animals. Embryos were collected by flushing the oviduct (1-3 days post coitum; dpc) or uterus (4-6 dpc) of donor females with pre-warmed medium (TCM-199+10% fetal bovine serum (FBS), Sigma). Where indicated, embryos were cultured *in vitro* in drops of RDH medium (RPMI:DMEM:Ham’s F10, Life Technologies, at 1:1:1) supplemented with 0.3% bovine serum albumin (Jin et al., 2000), under mineral oil, in a humidified incubator, at 38.5°C, 5% CO_2_ in air. Before *in vitro* culture, embryo coats were pre-digested by 45-s incubation in 0.5% pronase followed by 30 min incubation in TCM-199+10% FBS medium.

### Immunosurgery

Immunosurgery was performed to isolate the ICMs from rabbit blastocyst by removing the TE layer (Solter and Knowles, 1975). Prior to immunosurgery (IS), embryo coats were pre-digested as indicated earlier, and then removed mechanically. Immunosurgery was performed by 30 min. incubation in anti-rabbit serum (20% in TCM-199+10% FBS medium; 091M4751, Sigma), followed by 15-20 min. incubation in complement solution (20% in TCM-199+10% FBS medium; 234395, Millipore). Lysed cells were removed mechanically by pipetting.

### ICM culture

The isolated ICMs were cultured in the RDH medium supplemented with 0.3% bovine serum albumin, under mineral oil, in a humidified incubator, at 38.5°C, 5% CO_2_ in air (as indicated for whole embryos). ICMs were cultured under PrimoVision time-lapse system (Vitrolife) for brightfield time-lapse imaging for 24 or 48 hours.

### Immunostaining

Embryos and ICMs were fixed in 4% paraformaldehyde in PBS with 0.1% Tween-20 (Sigma) and 0.01% Triton X-100 (Sigma) for 20 min. at room temperature. Embryonic coats were removed mechanically after fixation (Püschel and Viebahn, 2010), unless otherwise indicated. Fixed embryos and ICMs were immunostained as previously described (Piliszek et al., 2017). Briefly, embryos were permeabilised in 0.55% Triton X-100 (Sigma) solution in PBS for 20 minutes, subjected to quenching unreacted aldehydes in NH4Cl for 10 minutes, and blocked in 10% donkey serum (Sigma) for 40 minutes. Primary antibodies were used at a 1:100, except anti-SOX2 antibody (abcam, 1:200). Antibodies are listed in Table 1. Embryos were incubated with primary antibodies overnight at 4°C. Alexa Fluor-conjugated secondary antibodies (Invitrogen: donkey anti-mouse Alexa 488, donkey anti-rabbit Alexa 647, donkey anti-goat Alexa 568) were used at 1:500. DNA was visualised using Hoechst 33342 (10 µM, Life Technologies, H3570).

**Table 1:** List of antibodies used.

| Target | Host | Company | Cat. Number |
| --- | --- | --- | --- |
| CDX2 | Mouse | BioGenex | MU3921_UC |
| GATA3 | Mouse | BioLegend | 653801 |
|  | Mouse | BioLegend | 653809 |
| OCT4 | Goat | R&D | AF1759 |
| SOX2 | Rabbit | Abcam | ab97959 |
|  | Goat | R&D | AF2018 |
| SOX17 | Goat | R&D | NL1924R |

### Imaging and image analysis

Embryos were placed on a glass-bottom dish (Thermo Fisher) and three-dimensional (3D) imaging was performed using an A1R Nikon inverted confocal microscope. Embryos were imaged in their entirety with a 20x objective using Z-stacks of 2 µm thickness.

Analysis of images was performed using IMARIS software (Bitplane AG). For cell number count, nuclei were identified using the ‘spot’ option with an estimated diameter of 7-10 μm. The total cell number was based on Hoechst chromatin staining, the number of cells positive for each transcription factor was based on appropriate immunofluorescent staining. The number of nuclei identified by IMARIS was confirmed manually. 3D confocal images were created by maximum intensity projection using the IMARIS ‘volume’ option.

### Embryo collection for gene expression analysis

*In-vivo*-obtained rabbit embryos were collected at successive developmental stages at 2 (morula), 3.0 (late morula), 3.25 (early blastocyst), 4, 5 and 6 dpc (blastocyst). The selected material was placed in a minimal volume of PBS in 1.5 ml tubes (low binding, Eppendorf), snap-frozen in liquid nitrogen and stored at −80°C.

### RNA extraction and cDNA synthesis

Total RNA was extracted with the High Pure miRNA Isolation Kit (Roche Diagnostics) following the manufacturer’s protocol, as previously described (Madeja et al., 2013; Piliszek et al., 2017). RNA quality and concentration were measured using NanoDrop c2000 (Thermo Scientific). For each sample, the reverse transcription reaction was performed from 100 ng of total RNA. cDNA synthesis was performed with the Transcriptor High Fidelity cDNA Synthesis Kit (Roche Diagnostics) following the manufacturer’s protocol. The samples were stored at −20°C.

### Quantitative real-time PCR reaction

Quantitative PCR (qPCR) was performed on a Roche Light Cycler 96 instrument. Calculations of gene expression level were based on the standard curve method (with series of 10 fold dilutions of known concentrations) with three reference genes: *H2AFZ*, *HPRT1* and *YWHAZ* (Mamo et al., 2007). The relative mRNA content was calculated to the normalised mean transcript level of the reference genes. Each sample was analysed in triplicate, with all of the primer sets chosen for the experiment. For each developmental stage, we collected six independent samples. The primer pairs were designed to span the introns. The reactions were carried out as previously described (Madeja et al., 2013; Piliszek et al., 2017). Product specificity was confirmed by melting-point analysis and agarose gel electrophoresis.

### Statistical analysis

Data processing and visualizations were done in a Jupyter notebook using the pandas 1.4.4, Matplotlib 3.7.2 and seaborn 0.12.2 Python libraries. The expression levels of *CDX2*, *GATA3*, and *OCT4* in rabbit embryos between consecutive stages of development were compared using the Kruskal-Wallis test (SciPy 1.10.1). Subsequent pairwise comparisons were conducted using the Conover test with a two stage FDR correction (scikit-posthocs 0.7.0).

## RESULTS

### Dynamic changes of embryo morphology during preimplantation rabbit development

We have previously established that a specific pattern of transcription factor expression in rabbit preimplantation embryos is not correlated to the time post coitum *per se*, but rather to the total cell number of the embryo (Piliszek et al., 2017) (Fig 1A). Our previous research also suggested that different blastocyst geometries are acquired during the consecutive stages of blastocyst maturation. To confirm that rabbit blastocyst indeed progresses through the specific geometries, and to gain more insight into the morphogenic processes driving this progression, in the current study we analysed the changes in the morphology of rabbit embryos during blastocyst cavity formation and expansion (Fig. 1B, Movie S1). To this end, *in vivo* fertilised embryos were collected at the late morula stage on day 3 of development, and time-lapse imaged during the 24-hour *in vitro* culture (n=16). We paid specific attention to the changes in shape and size of the blastocyst cavity, ICM and TE, in order to confirm the timely progression of the morphological landmarks (Fig. 1A, B, Movie S1). Our observations can be summarised as follows: initiation of cavity formation becomes first apparent as a U-shaped slit between the outer and the inner cells. The slit then expands into an ellipsoid cavity, with ICM located at one of the poles (stage VI). Next, in stage VII blastocysts, the cavity becomes more crescent-shaped, while multiple cytoplasmic bridges can be detected between TE and spherical ICM. At stage VIII, the cavity becomes nearly spherical, and ICM becomes more flattened against one pole of the cavity, although still clearly distinguished as a thickened/bulging area. At stage IX, the blastocyst cavity is spherical and ICM becomes flattened against polar TE. Later stages of rabbit blastocyst development maintain the same shape, and differ only in size, as blastocyst diameter further increases. During initial stages of cavitation (stage V, VI), TE cells are squamous, but later become flattened coincident with cavity expansion.

**Figure 1.**
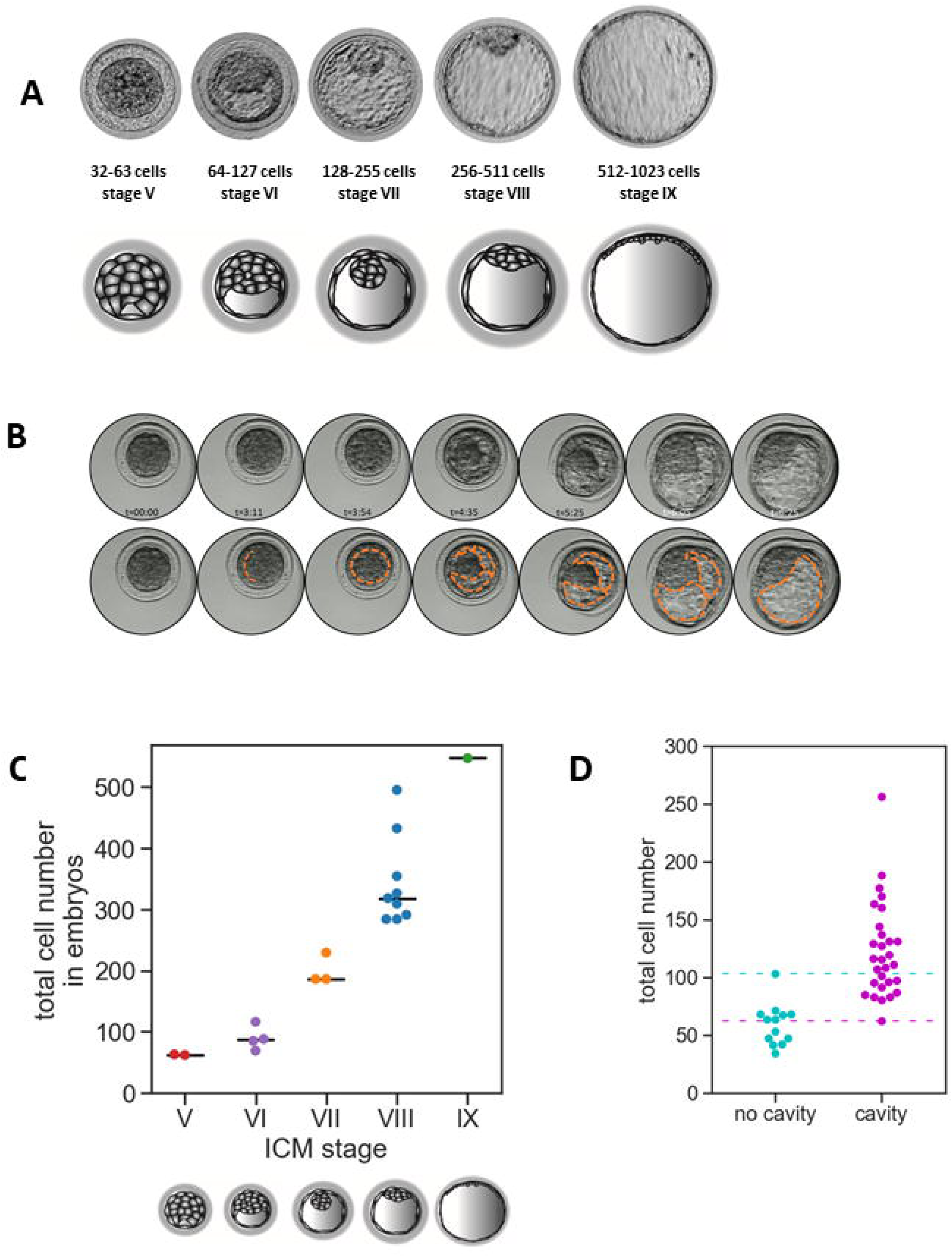
Dynamic changes of embryo morphology during rabbit blastocyst development. (A) Morphological landmarks are correlated with the cell number in E3.25 – E3.5 rabbit embryos *in vivo* (B) *In vitro* development of rabbit blastocyst. Selected sections from the Primo Vision time-lapse system. Orange dotted line denotes the boundaries of the blastocyst cavity (C) Rabbit embryo staging system based on cell number correlated with staging based on morphology. Calculation of cell number in embryos *in vivo* categorised to each stage based on the morphology. (D) Cell number in rabbit embryos at the time of cavitation. Dashed lines denote minimum and maximum cell number values for E2.75-E3.25 cavitated embryos.

To confirm our *in vitro* observations, we analysed a group of embryos of a certain morphology obtained *in vivo* and counted the cell number after fixation (Fig. 1C), which confirmed that the stage assessment based on embryo morphology closely correlated with cell number. Embryos obtained *in vivo* at embryonic day (E) 2.75-3.25, represented late morula and early blastocyst stage. Initial stages of the cavitation were observed at these timepoints in embryos having no fewer than 62 cells (Fig. 1D), which we further referred to as stage V blastocysts.

### CDX2 is a marker of mature TE in rabbit embryos

Having established the timing of events and morphological features related to a particular stage of the cavity formation and expansion, we sought to gain insight into the molecular players that are involved in TE formation in rabbits. qPCR analysis revealed that the *CDX2* transcripts are not expressed in rabbit until early blastocyst (E3.25), but are found at later stages (Fig. S1A).

In agreement with the qPCR data, the immunofluorescent analysis showed no CDX2 protein in rabbit morulae (n=28) and stage V and VI early blastocyst (Fig. 2). CDX2 was first observed in stage VII blastocysts (n=10), specifically in the TE cells, however, it was not initially detected throughout the whole TE. Instead, it was found only in single cells scattered within the TE (on average 9% of CDX2–positive cells per embryo in TE, but only 40% of embryos at this stage had CDX2-positive cells). The percentage of CDX2-positive cells in TE increased at the subsequent stages, reaching 80% by stage VIII (n=8) and 100% by stage X (n=5) (Fig. 2). The observed timing of CDX2 expression suggests that it becomes upregulated alongside the process of the cavity expansion, reaching ubiquitous expression in TE cells by stage IX (approximately E4.0). Therefore, unlike in the mouse, but in agreement with the data from human embryos (Niakkan and Eggan, 2013), CDX2 in the rabbit is a marker of mature TE.

**Figure 2.**
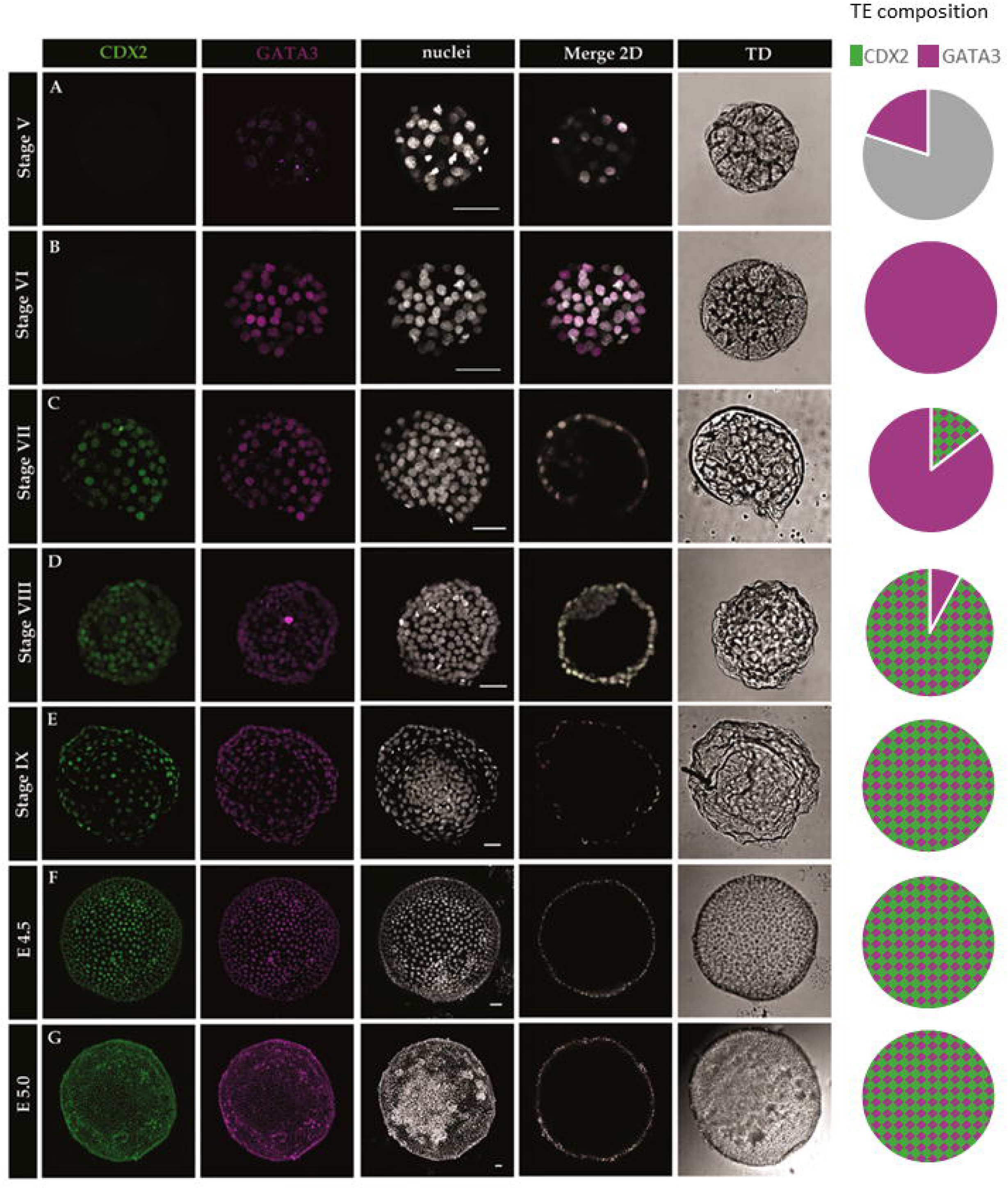
CDX2 and GATA3 are markers of TE in rabbit blastocyst. GATA3 is an early marker of TE in rabbit embryos, while CDX2 is a marker of mature TE. (A-G) Localisation of transcription factors CDX2 and GATA3 at consecutive stages of rabbit embryo development. GATA3 is first detected in stage V embryos (A), appearing in the majority of outside cells. CDX2 is first detected at stage VII (C), and initially appears only in a subset of TE cells. From stage IX (C, approximately E4.0), all TE cells are CDX2 and GATA3 double-positive. Each row represents a 3D reconstruction of a z-stack of a representative embryo (except merge 2D which is a single section). Green-CDX2; magenta-GATA3; white-nuclei (Hoechst); scale bar-50 um. Pie charts represent the percentage of GATA3 and CDX2 positive TE cells. Magenta-GATA3 only, Grey – double negative for GATA3 and CDX2; checkered magenta and green – GATA3 and CDX2 double positive.

### GATA3 is an early marker of rabbit trophectoderm

As CDX2 was not detected at the early stages of cavitation, we sought the additional factors that might drive the initial stages of TE specification in the rabbit embryos. GATA3 is another important transcription factor driving the TE differentiation in the mouse (Home et al., 2009; Ralston et al., 2010), and it has been also found in TE of other mammals, including humans (Gerri et al., 2020, Guo et al., 2020). Therefore, we analysed *GATA3* expression in rabbit embryos at E1.0-E6.0. The qPCR analysis revealed that *GATA3* is already expressed in E2.0 morulae, 24 hours ahead of cavitation, with the greatest transcript abundance at E3.25, early blastocyst stage (Fig. S1B). We found no GATA3 protein in rabbit morulae at E2.0 (n=6). We further analysed the localisation of GATA3 in E2.75-E6.0 rabbit embryos (Fig. 2). GATA3 was found in E2.75, fully compact late morula-stage embryos, just prior to cavitation (stage V morula) in 20% of the outside cells (n=4). In early cavitating blastocysts, 35% of outside cells were GATA3 positive (stage V blastocyst, around 60-70 cells). At subsequent stages, GATA3 was present in all the cells of the TE layer of rabbit E3.25-E6.0 blastocyst (stages VI and later) (n=18) (Fig. 2). In 50% of stage V, VI, VII blastocysts GATA3 was also present in single ICM cells located in the area adjacent to the blastocyst cavity (1-7 cells per embryo). By analysing the colocalisation of GATA3 and CDX2, we found that CDX2 localisation is always restricted to GATA3-positive cells (n=24) (Fig. 2).

In summary, the analysis of expression and localization of CDX2 and GATA3 in rabbit embryos reveals that both factors are specific markers of rabbit TE, but exhibit differences in the timing of expression. While GATA3 is an early marker that may play a role in the initial stages TE differentiation, CDX2 is a marker of the mature trophectoderm.

### OCT4 is expressed in both ICM and TE in rabbit blastocyst

Previous studies in mouse embryos suggested that CDX2 acts in ICM/TE specification by repressing activity of octamer-binding transcription factor 4 (OCT4, encoded by *Pou5f1/Oct-4*), a pluripotency factor which is downregulated in mouse TE at the early blastocyst stage (Niwa et al., 2005; Strumpf et al., 2005). However, it has been shown that in rabbit embryos OCT4 persists in TE until the mid-blastocyst stage (Chen et al., 2012b; Kobolak et al., 2009). To verify these findings, we analysed *OCT4* transcript expression by qPCR in E2.0-E6.0 rabbit *in vivo* embryos. We found that *OCT4* was expressed at all of the analysed stages, however the mRNA expression significantly increased at E3.0 late morula stage (*P*<0.05) and decreased in E6.0 late blastocysts (*P*<0.005) (Fig. S1C). We further analysed the distribution of OCT4 protein in stage V to IX rabbit blastocysts. OCT4 protein was present in rabbit embryos at all stages analysed and detected in nuclei of all cells in both TE and ICM (Fig. 3). This result suggests that in the rabbit embryos downregulation of OCT4 is not a prerequisite for TE specification nor differentiation.

**Figure 3.**
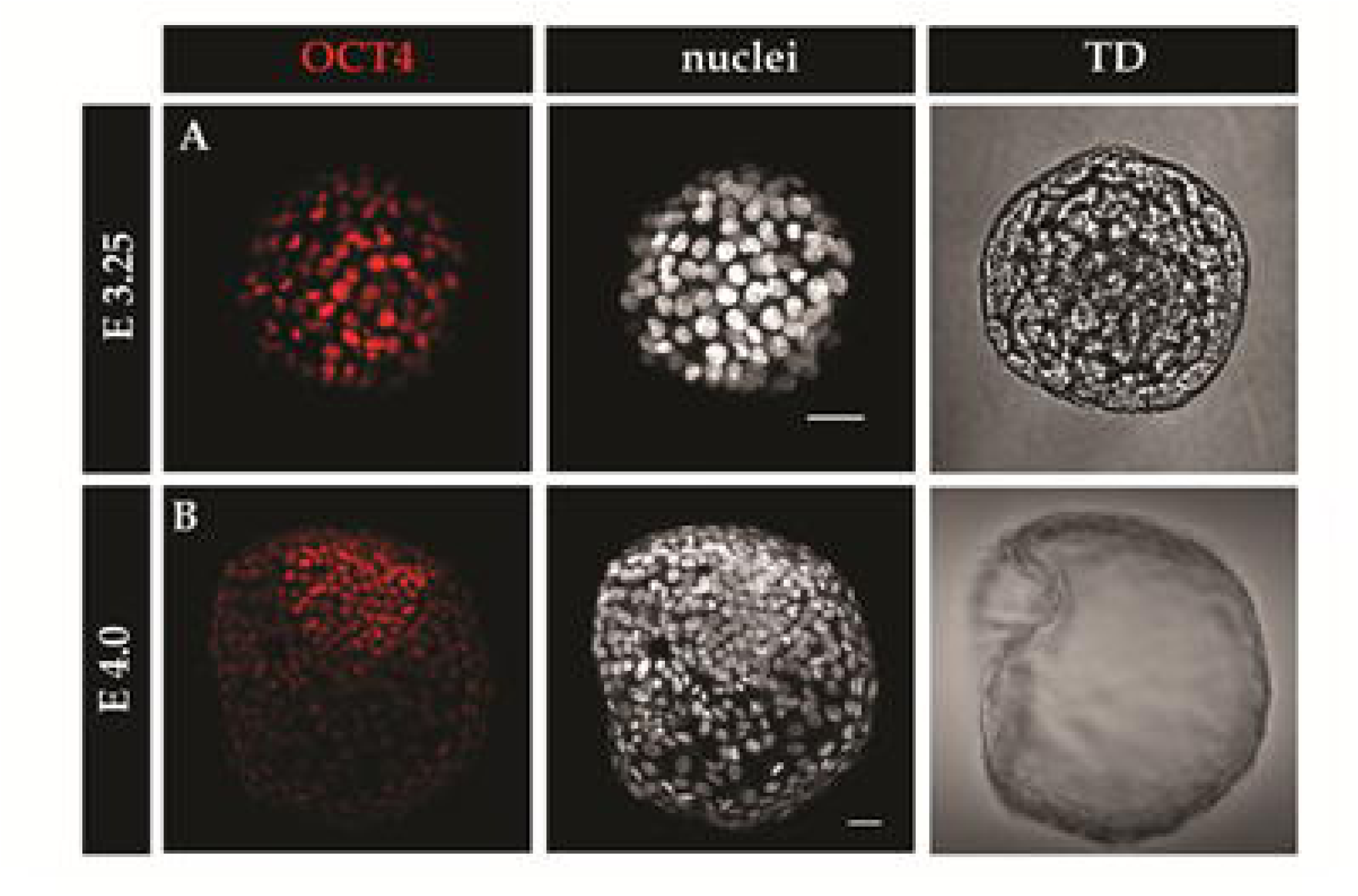
OCT4 is expressed in both TE and ICM in rabbit blastocyst. (A) Localisation of OCT4 in embryos at E3.25 and E4.0. Red – OCT4; white – nuclei (Hoechst); TD – transmitted light; scale bar – 50 µm.

### Rabbit mid-blastocyst ICMs are able to restore TE layer

As OCT4 is retained in rabbit TE for much longer than in mouse embryo, and its presence does not seem to be detrimental for the initial stages of TE maturation, we were wondering what are the functional consequences of such prolonged OCT4 presence in TE. We hypothesised that isolated rabbit ICMs from later blastocyst stages might be still able to re-create TE layer, as the high levels of OCT4 will not prevent TE differentiation.

In order to investigate the ability of ICM to restore TE, we immunosurgically removed the TE layer from rabbit blastocysts, and followed the development of isolated ICMs cultured *in vitro* (further called IC-ICMs – isolated and cultured ICMs) (experimental design – Fig. 4). As we previously established that the cavitation timing does not depend directly on the time *post coitum* in rabbits, we staged IC-ICMs according to the geometry of blastocyst they were derived from, using information about geometries progression (Fig. 1A) correlated with our previously developed staging system (Fig. 1A). To this end, E3.25 and E3.5 blastocysts were assigned into five groups, according to the stage of rabbit blastocyst development: V, VI, VII, VIII and IX (Fig. 1A). Isolated ICMs (n=147) were subsequently cultured for 24 or 48 hours under PrimoVision time-lapse imaging system.

**Figure 4.**
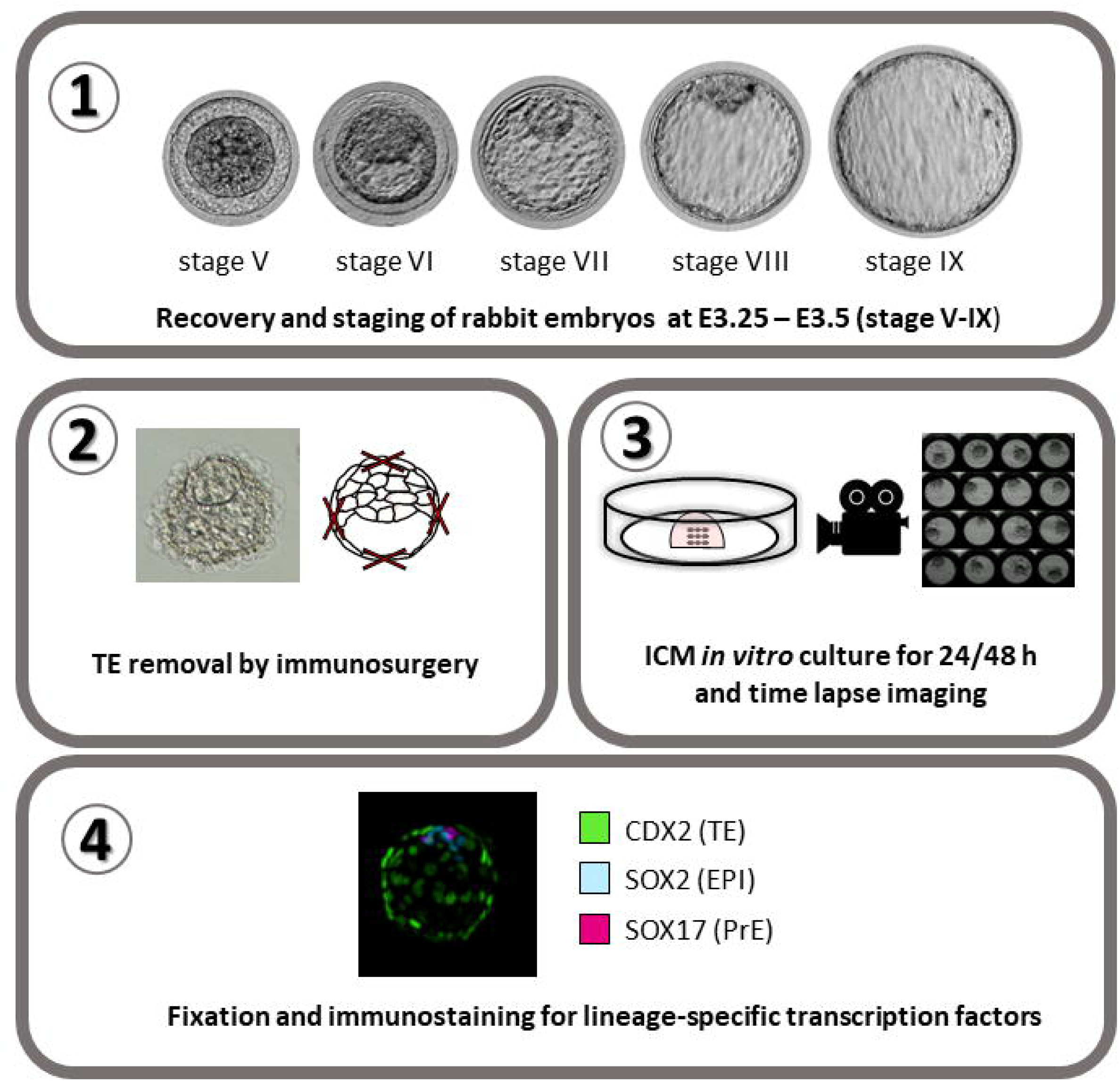
Challenging ICM potency – experimental design. (1) Embryos were first selected according to the morphology and then (2) processed through immunosurgery (IS) to remove TE later. Next, (3) isolated ICMs were cultured under PrimoVision time-lapse imaging system for 24 or 48h and then (4) fixed, immunostained for lineage-specific markers and imaged under the confocal microscope.

Following 48h *in vitro* culture, we observed three distinct geometries of IC-ICMs. Type A IC-ICMs formed a blastocyst-like structure, restoring TE layer in the process similar to blastocyst cavitation *in vivo,* progressing through the same morphological landmarks as cavitating intact rabbit blastocysts (Fig. 5A, Movie S2). Type B developed a small, halo-like cavity around a centrally localised group of cells (Fig. 5B, Movie S3). Type C did not restore any cavity and remained as a compact cluster of cells (Fig. 5C). ICM stage-of-origin strongly impacted IC-ICM development (Fig. 5D). Stage V IC-ICMs restored type A – blastocyst-like geometry in 100% of the cases (n=23). The potential to restore blastocyst-like structure decreased with developmental timing progression: Type A geometry was observed in 66.7% of stage VI IC-ICMs, 45.2% of stage VII, 15% of stage VIII and no type A geometry was re-formed from stage IX IC-ICMs. Concomitantly, halo-like cavity (type B) was not observed in IC-ICMs from stage V, but it was found with an increasing incidence at later stages – 4.8% in stage VI, 35.5% in stage VII, 55.5% in stage VIII, and 100% IC-ICMs from stage IX restored halo-like cavity (n=8).

**Figure 5.**
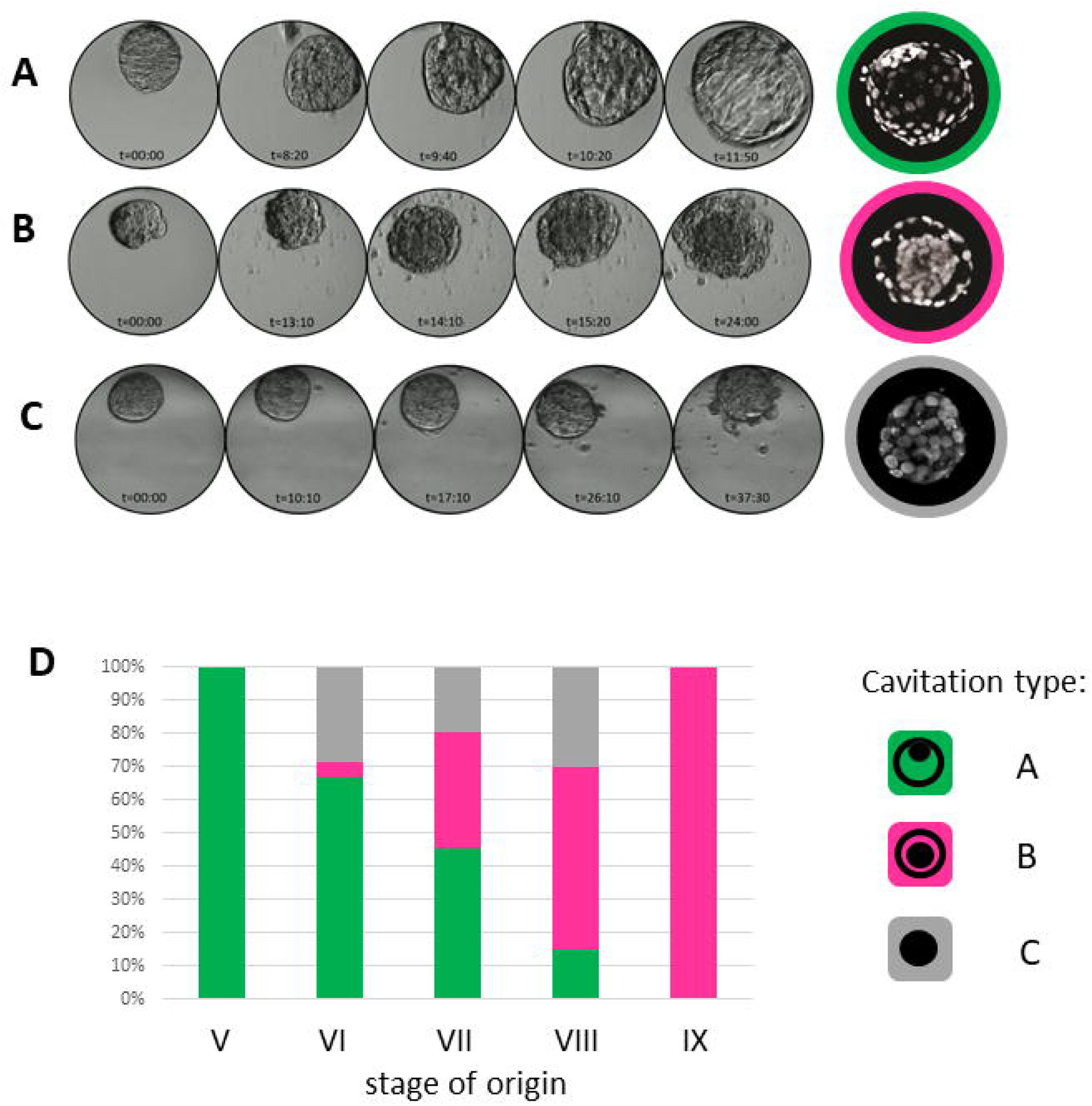
In vitro development of isolated rabbit ICMs. Time-lapse images of development of isolated rabbit ICMs during 48 h *in vitro* culture, followed by a corresponding cross-section of chromatin-stained IC-ICM (white, Hoechst) (A) Type A “blastocyst” cavitation. IC-ICMs isolated from blastocyst stages V, VI, VII follow a similar pattern of cavitation to intact rabbit embryos (compare to Figure 1) (B) Type B “halo” cavitation. IC-ICMs isolated from blastocyst stages VIII and IX fail to develop a blastocyst-like cavity but instead form a ring-shaped cavity around central cell mass. (C) Type C “no cavitation”. Despite *in vitro* culture, some IC-ICMs fail to re-cavitate (D) Percentage contribution of each cavitation type depending on the stage of embryo origin. Green-Type A, Magenta-Type B, Grey – Type C

### CDX2 and GATA3 expression coincides with restoration of functional TE-layer in recavitated IC-ICMs

IC-ICMs (n= 98) were fixed and stained for lineage markers: SOX2 for Epi, SOX17 for PrE and CDX2 for TE (Fig. 6A). We noticed that the number of CDX2+ cells decreases with the stage of isolated IC-ICM i.e. IC-ICMs isolated from later stages (VII, VIII) have fewer CDX2+ cells (fewer recavitated IC-ICMs with CDX2+ cells and also fewer CDX2+ cells per each recavitated IC-ICM). In ICMs isolated from stage IX, we did not find any CDX2+ cells, suggesting that at this stage all ICM cells lost their ability to differentiate towards TE (Fig. 7).

**Figure 6.**
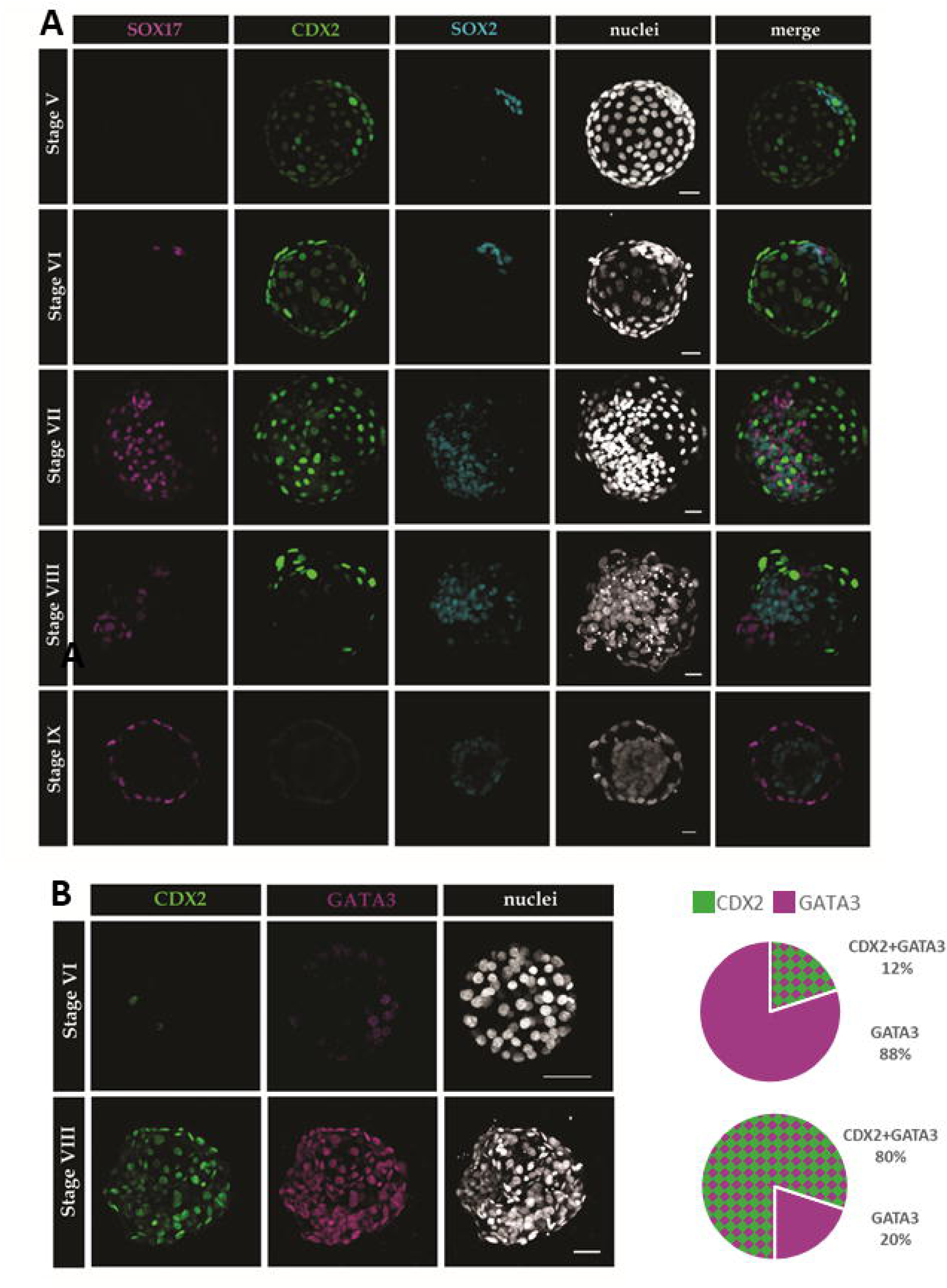
**(A) Rabbit ICMs retain full potency until mid-blastocyst stage** Localisation of PrE (SOX17), TE (CDX2) and EPI (SOX2) markers in IC-ICMs after 48h *in vitro* culture. In IC-ICMs obtained from stages V and VI, CDX2 and SOX2 are found in many cells, outside and inside, respectively and SOX17 was found in single cells only. In IC-ICMs isolated from stage VII and VIII, all three transcription factors are observed widely spread. IC-ICMs originating from stage IX comprise SOX17 and SOX2 positive cells, but CDX2 is not observed. Each row represents a 3D reconstruction of a z-stack of a representative embryo. Magenta – SOX17; Green – CDX2; Blue – SOX2, white – nuclei (Hoechst); scale bar – 30 µm. **(B) Restored TE of IC-ICMs co-expresses TE markers GATA3 and CDX2.** Localization of CDX2 and GATA3 in IC-ICMs after 48h of *in vitro* culture. In IC-ICMs originating from stage VI, GATA3 is observed in a majority of outside cells, however CDX2 is found only in single cells. In IC-ICMs isolated from stage VIII, both TE markers are mostly colocalised, however higher contribution of GATA3 positive cells is still notable in the newly reformed TE. Each row represents a 3D reconstruction of a z-stack of a representative embryo. Green-CDX2; magenta-GATA3; white-nuclei (Hoechst); scalebar-50 µm. Pie charts represent the percentage of GATA3 and CDX2 positive TE cells. Magenta-GATA3 only, checkered magenta and green – GATA3 and CDX2 double positive.

**Figure 7.**
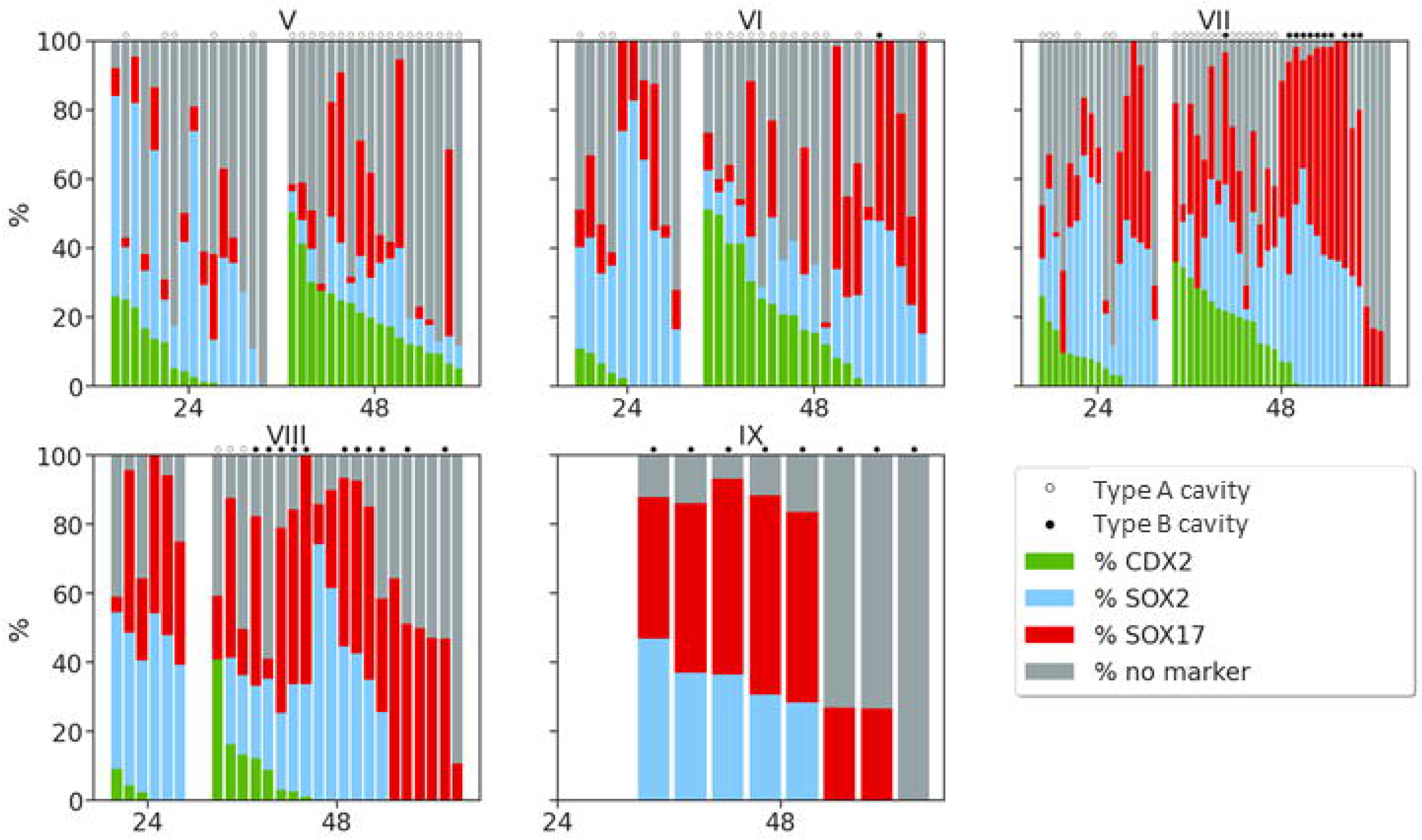
Contribution of CDX2, SOX2, SOX17 positive, and triple-negative cells in all IC-ICMs isolated from stage V, VI, VII, VIII and IX blastocysts, after 24 and 48 h of *in vitro* culture. IC-ICMs with type A cavitation are marked with“°”, type B cavitation – with”•” Each column represents a single IC-ICM, data are presented in order of the decreasing CDX2 contribution. All three transcription factors were observed in IC-ICM after culture, except in IC-ICMs obtained from stage IX, where CDX2 was not observed. Green – CDX2; Blue – SOX2; Red – SOX17; Grey – triple-negative cells

In the rabbit blastocyst, GATA3 is an early TE marker that precedes CDX2 expression. To further uncover the extent of TE specification in recavitated IC-ICMs, we analysed GATA3 distribution. After 48h of culture, GATA3 was present in outside cells of 70% of recavitated IC-ICMs (n=20) originating from V-IX stage blastocysts. Re-formed TE of type A IC-ICMs contained mostly CDX2^+^ GATA3^+^ double-positive cells, although in 14.2% of embryos we also found GATA3+, CDX2-cells (n=14), which resembles the expression pattern in early *in vivo* blastocyst. In IC-ICMs isolated from stage VI blastocysts, only 12% of outside cells were GATA3+ CDX2+ double-positive, while in those isolated from stage VIII blastocysts up to 80% were double-positive (Fig. 6B). As in intact embryos, we found no CDX2+ GATA3-cells in any of the IC-ICMs.

### TE specification is initiated in rabbit ICM shortly after isolation

To better understand the dynamics of IC-ICM development and differentiation, we additionally analysed stage V-VIII IC-ICMs after a shorter, 24-hour *in vitro* culture (n=49) (Fig. 7). We found that IC-ICMs isolated from stage V restored type A cavity in 31.3% of the cases, from stage VI in 40% and from stage VII in 41% of the cases, but no cavity was formed after 24h culture of IC-ICMs from stage VIII. This indicates that TE re-formation is a relatively rapid process, as we observed type A cavitation 24h after ICM isolation, and in some cases already after 10h of *in vitro* culture. In contrast, type B cavitation was never observed after 24 hours of IC-ICM *in vitro* culture.

### IC-ICMs express markers of TE, Epi and PrE lineages

Having established that isolated rabbit ICMs are able to restore a blastocyst-like structure, which has an outside GATA3+, CDX2+ TE layer, we asked whether the inner cells of the IC-ICM also differentiate into appropriate lineages. In order to do it, we tested IC-ICMs for the presence of PrE (SOX17) and Epi (SOX2) lineage markers (Fig. 6 and 7). The majority of IC-ICMs contained both SOX2+ and SOX17+ cells, and double-positive cells were never present. Epiblast marker SOX2 was present in 86.5% of all recavitated IC-ICMs (n=96). The percentage of SOX2+ cells increased in IC-ICMs derived from later stages – namely, IC-ICMs from stage V contained on average 12.3% of SOX2+ cells, from stage VI – 16.9%, from stage VII – 23.9%, from stage VIII – 22.4% and from stage IX – 25.2%. In all of these cases, SOX2+ cells were present in the inside compartment, and in 39.6% IC-ICMs a small number of SOX2+ cells was also found in the outside compartment (1-5 cells per embryo) (Fig. 8). PrE marker SOX17 was present in 89% of re-IC-ICMs (n=96). The overall percentage of SOX17+ cells was higher in the IC-ICMs derived from later stages – namely, IC-ICMs from stage V contained 22% SOX17+ cells, from stage VI – 22%, from stage VII – 32%, from stage VIII – 40.4% and from stage IX – 42.7%. In 84.4% of IC-ICMs SOX17 was found in the inside localised cells, and in 57.3% in the outside cells (Fig. 8).

**Figure 8.**
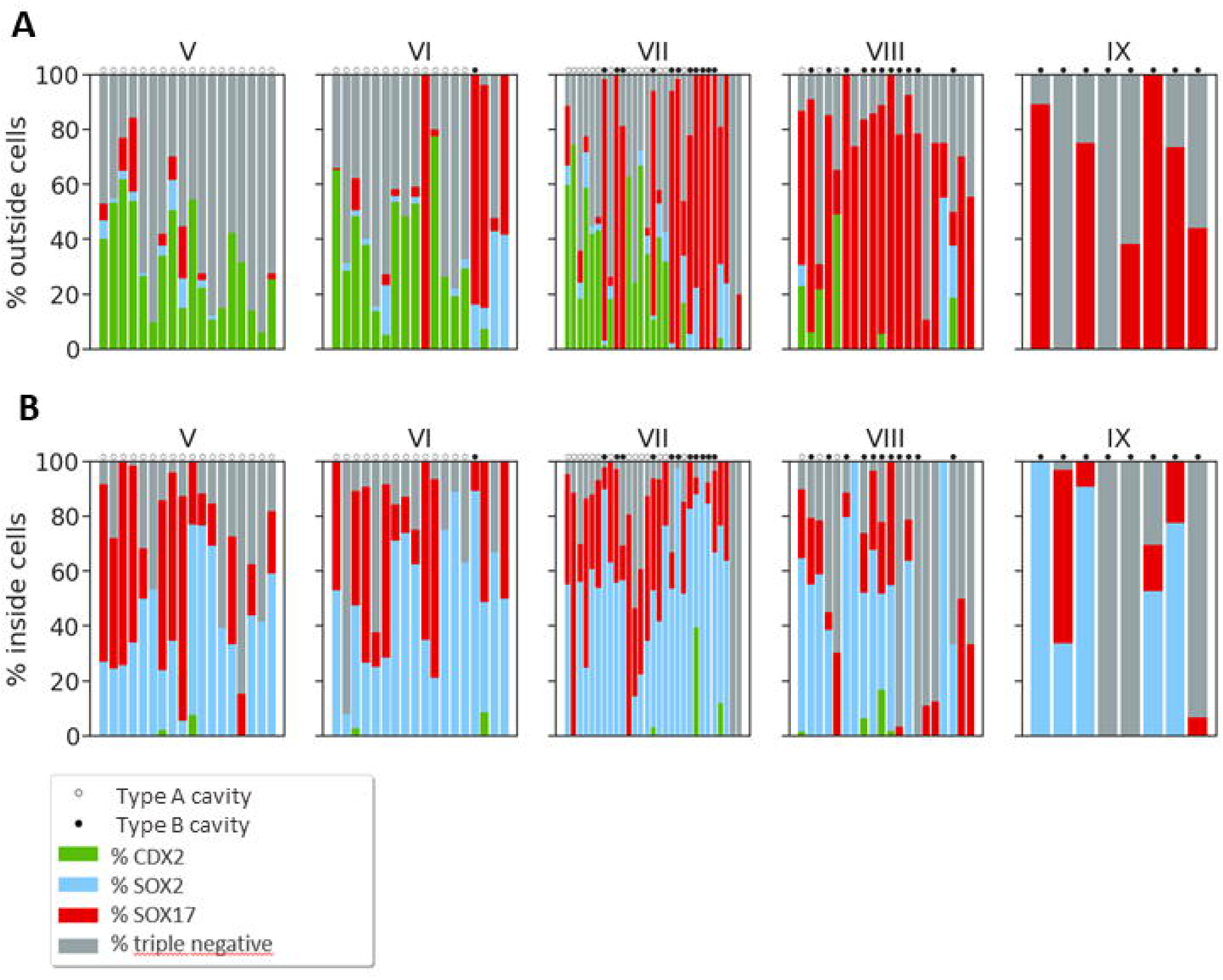
Percentage contribution and differentiation of inside and outside cells in all IC-ICMs. isolated from stage V, VI, VII, VIII and IX blastocysts after 24 and 48 h *in vitro* culture. (A) CDX2, SOX2, SOX17 and triple-negative cells in re-IC-ICMs in outside cells. (B) CDX2, SOX2, SOX17 and triple-negative cells in re-IC-ICMs in inside cells. IC-ICMs with type A cavitation are marked with “o”, type B cavitation – with”•”. Each column represents a single IC-ICM, data are presented in order of the decreasing cell count in the embryo.

We also analysed the correlation between the number of cells positive for each single lineage marker and the number of triple-negative cells in each embryo (Fig. S2). Analysis for SOX2^+^ and CDX2^+^ cells did not show any correlation, however, we noticed that the number of SOX17^+^ cells is negatively correlated with the number of triple-negative cells – we observed more SOX17^+^ cells in re-IC-ICMs with a small percentage of triple-negative cells. SOX17+ cells were never found in IC-ICMs with more than 80% of triple-negative cells, and the IC-ICMs with less than 20% of triple-negative cells often had low or no contribution of CDX2+ cells.

### Rabbit IC-ICMs isolated from late blastocyst form SOX17+ outside layer

Next, we analysed whether each of the observed cavitation geometries (type A, B or C) are correlated with a specific expression pattern of lineage-specific transcription factors in IC-ICMs following 48-hour *in vitro* culture (Fig. 7 and 8).

Analysing the distribution of the TE, PrE and Epi markers in the IC-ICMs after 48h culture we noticed that CDX2 and SOX17-positive cells were present in both inside and outside locations. We noted that the contribution of SOX17+ cells to the outside compartment was higher in the IC-ICMs originating from later-stage rabbit blastocysts. SOX17+ cells constituted only 5.3% of outside cells in IC-ICMs derived from stage V, 12% from stage VI, 28.1% from stage VII, 55.5% from stage VIII and 62.8% from stage IX. Overall, in 41% of IC-ICMs SOX17+ cells were present in the outside layer only. To quantify that further, we analysed the distribution of cells within the re-IC-ICMs in the inside versus outside compartments after 48 hours of culture in relation to the type of cavity restoration geometry (Fig. 8).

CDX2-positive cells were present in nearly all of the type A IC-ICMs from all stages (48 out of 49 IC-ICMs), and in all of these cases CDX2 was predominantly present in the outside cells. Only 2 IC-ICMs from stage V, 1 IC-ICM from stage VI and 1 from stage VIII contained 1-3 CDX2-positive cells in the inside compartment (Fig. 8 A, B).

The blastocyst-like structures (type A) contained all three cell lineages, properly sorted into their respective compartments (Fig. 6). In addition to CDX2 and GATA3 positive outside TE layer, type A IC-ICMs re-formed an ICM which contained SOX2+ cells (in 91% of the cases) and SOX17+ cells (85% of the cases) (Fig. 8 B). In none of the cases the analysed markers were colocalised, suggesting that rabbit IC-ICMs are capable of proper differentiation and segregation into the first three cell lineages within 48h period.

In type B IC-ICMs after 48h culture, the percentage of CDX2-positive cells almost never exceeded 15% per ICM (in 30 out of 31 IC-ICMs). No CDX2-positive cells were present in type B IC-ICMs from stages VI and IX (Fig. 7), whereas 45% of stage VII and 18% of stage VIII IC-ICMs contained some CDX2 positive cells, mostly located in the inside compartment. The outside layer of type B IC-ICMs was largely devoid of CDX2 (no CDX2-positive cells in 26 out of 31 type B IC-ICMs, and only up to 5 cells in the remaining 5).

Careful analysis of composition of the outside layer in relation to the type of cavitation geometry revealed that while in 95.9% of type A IC-ICMs outside layer contained CDX2+ cells, in 93.6% of type B IC-ICMs the outside layer was composed mostly of SOX17+ cells (Fig. 8 A).

Our data indicate that rabbit ICMs from earlier stages have a higher capability to restore the TE layer. This ability is gradually lost, although still maintained in some ICMs from stage VIII. Later stage IC-ICMs form an outside layer of SOX17-positive cells and are only able to restore the halo-like cavity. In summary, the ability to differentiate towards TE, defined by both type A blastocyst-like cavity re-formation and expression of CDX2 and GATA3 in newly formed TE, is gradually lost, giving way to the formation of SOX17-positive outside layer and type B ‘halo’ cavity geometry.

## DISCUSSION

Recent studies suggest the existence of substantial differences in formation of the first cell lineages between mammals (reviewed in (Plusa and Piliszek, 2020)). However, the significance and origin of these differences are not yet fully understood. In this work we investigate how the cellular plasticity of early mammalian embryos relates to developmental time scale and changes in gene expression using rabbit isolated ICMs.

In the rabbit embryo, the morphological distinction between the TE and the ICM becomes first apparent around E3.0, in embryos composed of around 64 cells. Observation of *in vivo* obtained embryos, as well as time-lapse imaging of embryos collected at the late morula stage and subsequently cultured *in vitro*, revealed distinct features of early rabbit blastocyst morphology and cavity expansion. Indeed, we observed several differences between the cavity formation of mouse versus rabbit embryos. In the rabbit, we have not observed multiple mini-cavities, and consequently, no merging thereof. Instead, initiation of cavity formation appears in a form of a U-shaped slit between the outer layer of the morula and its inner cells. Unlike in the mouse embryos (Ryan et al., 2019), cells of this outer layer (i.e. trophectoderm) are not flattened or stretched, but maintain the round shape, similarly to early bovine TE cells (Maddox-Hyttel et al., 2003).

CDX2 and GATA3 are the main drivers of TE fate in the mouse (Home et al., 2009; Ralston et al., 2010; Strumpf et al., 2005). Their TE-specific localisation has been also confirmed in a number of other mammalian species (reviewed in (Plusa and Piliszek, 2020)). Previous studies suggested a late onset of CDX2 expression in rabbit embryos (Chen et al., 2012a), as CDX2-positive cells were found only in a subset of E4.0 early blastocyst TE. In agreement with this study, we found no CDX2-positive cells in E3.0 morulae prior to cavitation. Moreover, a more detailed analysis of rabbit embryos between E3.0 and. E4.0 allowed us to establish that CDX2 becomes first apparent in stage VII blastocyst. Interestingly, it can be initially detected only in single cells and becomes expressed ubiquitously throughout TE only at stage IX (approximately E4.0). This analysis shows that cavitation in rabbit embryos is executed in the absence of CDX2, and therefore *CXD2* expression is not a prerequisite for the initial stages of TE formation. A similar observation has been made in pig and human embryos, where the initiation of the TE formation takes place before the CDX2 becomes expressed (Bou et al., 2017; Liu et al., 2015; Niakan and Eggan, 2013). Indeed, in bovine and porcine embryos, as well as in the mouse, downregulation of *CDX2* does not disrupt cavitation and early stages of TE formation (Berg et al., 2011; Goissis and Cibelli, 2014; Sakurai et al., 2016; Strumpf et al., 2005; Wu et al., 2010). However, CDX2 is necessary for the later stages of TE differentiation, as loss of CDX2 results in failure in correct TE proliferation (cow: (Berg et al., 2011)), maintenance of epithelial integrity (mouse: (Strumpf et al., 2005; Wu et al., 2010)) and cell polarity (pig: (Bou et al., 2017)) which ultimately leads to failure in embryo implantation. Despite the lack of clear requirement of early *Cdx2* expression for TE cell fate specification, it is expressed in mouse morulae as early as the 8-cell stage (Dietrich and Hiiragi, 2007; Strumpf et al., 2005). This disparity may be explained by possible differential regulation of pre- and post-cavitation *Cdx2* expression. While the latter is dependent on the Hippo pathway (Hirate et al., 2013; Nishioka et al., 2008, 2009; Sasaki, 2017), the early, morula stage expression is regulated by Notch signalling in the mouse (Menchero et al., 2019; Rayon et al., 2014) and might be entirely missing in the other mammalian species.

In search of a factor capable of initiation of TE program, and a suitable early marker of TE lineage, we investigated the distribution of GATA3, a transcription factor previously reported in mouse (Home et al., 2009; Ralston et al., 2010), which has been also recently shown to be a TE-specific marker in bovine and human embryos (Gerri et al., 2020; Guo et al., 2021). Immunofluorescent analysis of rabbit preimplantation embryos showed that GATA3 is an early, specific and robust marker of rabbit TE (Fig. 2). It is first detected in outside cells of rabbit stage V late compact morula (at approximately E3.0), and throughout subsequent development it is present specifically in all cells of rabbit TE. The co-localization analysis also showed that CDX2 protein expression is restricted only to GATA3-positive cells (although up to stage VIII, GATA3-positive, CDX2-negative cells are present), which suggests that GATA3 might be a prerequisite for CDX2 expression. Additionally, we detected *GATA3* transcripts 24h prior to protein, which may suggest that the TE program is initiated in the rabbit embryo as early as E2.0, or that it might require an additional trigger just before cavitation.

Analysis of *OCT4* mRNA expression in rabbit embryos *in vivo* revealed significant increase in transcript levels at E3.0 late morula stage. Consecutively, at E3.25 early blastocyst stage, *OCT4* became downregulated (P<0.05), while *GATA3* transcript levels significantly increased (P<0.05). However, at this stage we have not detected *CDX2* transcripts. Analysis of OCT4 protein localisation in *in vivo* rabbit blastocyst confirmed its presence in both ICM and TE, in agreement with previous reports (Chen et al., 2012b; Kobolak et al., 2009). This indicates that in rabbit OCT4 downregulation is not necessary to initiate or maintain the TE program up to the mid-blastocyst stage, and further supports inter-species differences in gene regulatory networks, underlying differences in the mechanisms of lineage specification. Indeed, OCT4 has been detected in both ICM and TE in a number of species, including human, cattle and pig (Berg et al., 2011; Hansis et al., 2000). Although in the mouse embryo CDX2 acts as a direct transcriptional repressor of *Oct-4* (Huang et al., 2017; Ralston and Rossant, 2008), it is not the case in porcine embryos (Bou et al., 2016), where CDX2 has been shown to promote OCT4 proteasomal degradation, but not transcriptional control. Comparative studies of bovine and mouse embryos revealed the existence of a cis-acting regulatory region required to suppress TE-specific *POU5F1* transcription in murine embryos (Berg et al., 2011). This species-specific enhancer have been found in mice, but not in cattle, humans and rabbits (Berg et al., 2011), clearly confirming differences in gene regulatory networks governing early lineage specification between mammals.

Next, we sought to analyse the differentiation potential of rabbit embryos with respect to TE versus ICM specification. In the mouse embryo, both TE cells and ICM cells become mostly restricted in their developmental potential soon after cavitation (Gardner, 1983; Nichols and Gardner, 1984; Posfai et al., 2021; Suwińska et al., 2008). However, isolated cells of morphologically distinguishable TE are able to contribute to ICM lineages in chimaera studies of bovine and human embryos (Berg et al., 2011; De Paepe et al., 2013). Recent studies have also shown that in bovine embryos ICM potential is maintained for a longer period, as isolated ICMs of E6.0 blastocysts are able to regenerate TE layer (Kohri et al., 2019). This raises an interesting question of inter-specific differences of ICM/TE differentiation mechanism. An early restriction of cell fate might be a direct result of unusually rapid development of mouse embryo, which is ready for implantation in just 4 days. In other mammalian species, the preimplantation period is extended over several days or even weeks, which may not require such immediate specification decisions (reviewed in (Piliszek and Madeja, 2018), and may result in a longer period of plasticity or even totipotency.

Our analysis of the development of isolated rabbit ICMs uncovered that after 48 h culture they were able to form two distinct types of cavitated structure. Type A had a typical blastocyst morphology, with a single large cavity and ICM located peripherally within it, type B formed a halo-like cavity around a centrally located mass of cells. We also observed that the morphological changes during type A cavity re-formation progressed through the same stages as in normal blastocyst development (with respect to the order of events, shape of the cavity and shape of the outer layer of cells). IC-ICMs potential to form each type of cavity depended on the stage the ICM was derived from, with earlier stages (V, VI) preferentially forming type A, and later stages (VIII, IX) preferentially forming type B cavity (Fig. 5). Moreover, we noted that type A cavitation is a relatively more rapid process, as after 24h *in vitro* culture we already observed type A, but no type B morphology. Immunofluorescent analysis revealed that type A structures contained cells expressing markers of PrE (SOX17), Epi (SOX2) and TE (CDX2 and GATA3) localized exclusively in the appropriate compartment. This allows us to conclude that stage V and VI IC-ICMs are not restricted to EPI or PrE cell fate but retain the potential to differentiate into all three lineages of the mammalian blastocyst, including TE. However, the ability for type A cavity regeneration is progressively diminished in the ICMs isolated from later stages and entirely lost by stage IX (approximately E4.0 expanded blastocyst). Moreover, we have noted that in type B IC-ICMs the outer layer is composed of CDX2-negative, SOX17-positive cells. Our previous studies showed that stage IX ICM is entirely composed of already sorted EPI and PrE compartments (Piliszek et al., 2017), therefore this loss of plasticity to regenerate TE coincides with possible differentiation of the entirety of ICM cells to the other two lineages. Mouse isolated ICMs lose the re-cavitation potential already at the early blastocyst stage (Gardner, 1983; Nichols and Gardner, 1984), but similarly to rabbit embryos, later ICMs form an outer layer of PrE, GATA4-positive cells, even though a halo-like cavity was not observed in this case (Wigger et al., 2017). A recent analysis of human ICM explants revealed their potency to generate TE cells *in vitro*, but this ability was later lost to allow for amnion cells differentiation (Guo et al., 2021).

In summary, this body of evidence suggests that the TE specification program potentially employs the same set of lineage-specific transcription factors across the eutherian mammals. Importantly, loss of developmental plasticity and disparity of timing in ICM versus TE differentiation reflect inter-species differences, coincident with variances in species anatomy and reproductive physiology. The precise length of the TE-permissive period varies between species and likely depends on the length of the preimplantation period (4 days in mouse to over 20 days in cow) (Finn and McLaren, 1967; Wathes and Wooding, 1980), and more specifically on the timing of differentiation of other extraembryonic lineages. Taken together with recent findings in other species, our data indicates that mammalian ICM cells have time-limited potential to regenerate TE, but this ability has to be gradually lost to allow for differentiation into another extraembryonic epithelial layer.

## Acknowledgments

We thank technical staff of IGAB PAS for animal husbandry and help with embryo derivation.

## Supplementary files

**Figure S1**

Expression levels of (A) *CDX2*, (B) *GATA3* and (C) *OCT4* mRNA in rabbit *in vivo* embryos at consecutive stages of development (E1.0 to E6.0). Differences in expression levels between consecutive developmental stages were tested using the Kruskal-Wallis test and subsequent pairwise comparisons were conducted using the Conover test with a two stage FDR correction (* for p<0.05, ** for p<0.005). Error bars represent standard error of the mean (SEM).

**Figure S2**

Correlation between transcription factor expression (SOX2, SOX17, CDX2, and ‘no marker’ triple-negative cells) and cell localisation (all cells (A), inside (B) outside (C)) in IC-ICMs after *in vitro* culture

**Movie S1**

***In vitro* development of rabbit blastocyst.** Time-lapse bright field movie of a rabbit embryo, fertilised *in vivo*, recovered at E3.0 late morula stage, and subsequently cultured *in vitro* under PrimoVision system.

**Movie S2**

**Type A “blastocyst” IC-ICM cavitation**. Time-lapse bright field movie of a rabbit IC-ICM cultured *in vitro* under PrimoVision system, developing a type A cavity geometry.

**Movie S3**

**Type B “halo” IC-ICM cavitation**. Time-lapse bright field movie of a rabbit IC-ICM cultured *in vitro* under PrimoVision system, developing a type B cavity geometry.

