## Supplementary figures and images for "Stage-sensitive potential of isolated rabbit ICM to differentiate into extraembryonic lineages"

### Figure S1

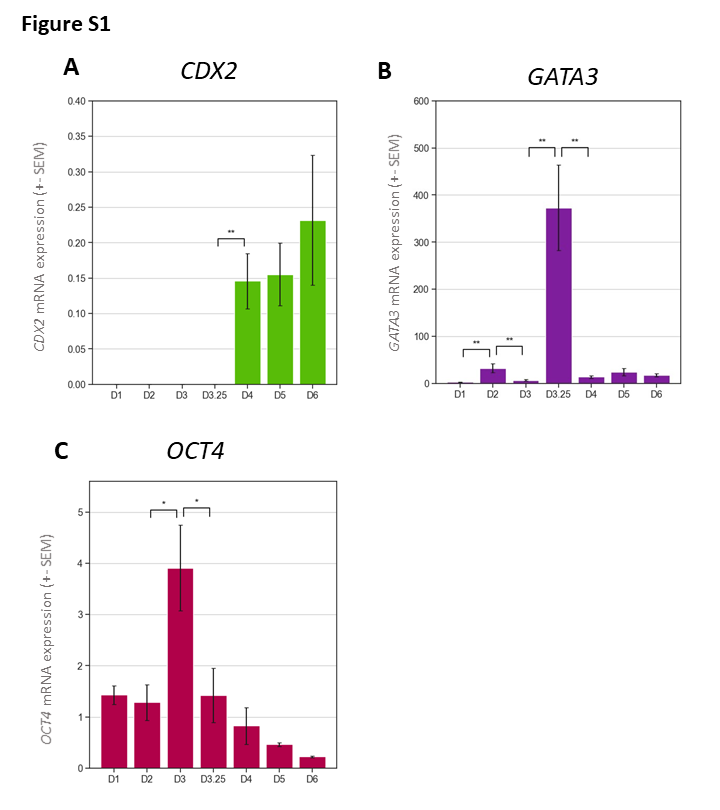

### Figure S2

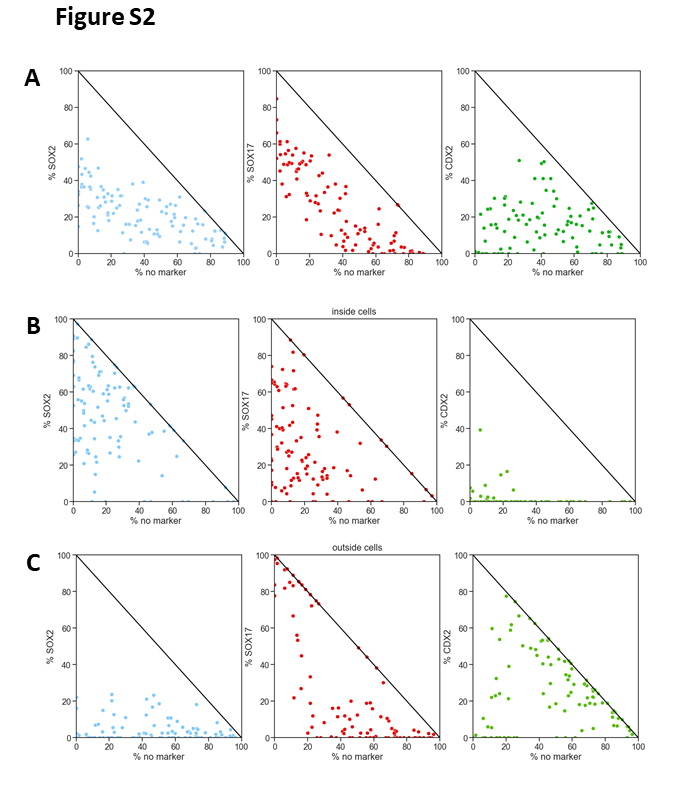
